# SpaCCC: Large language model-based cell-cell communication inference for spatially resolved transcriptomic data

**DOI:** 10.1101/2024.02.21.581369

**Authors:** Boya Ji, Liwen Xu, Shaoliang Peng

## Abstract

Drawing parallels between linguistic constructs and cellular biology, large language models (LLMs) have achieved remarkable success in diverse downstream applications for single-cell data analysis. However, to date, it still lacks methods to take advantage of LLMs to infer ligand-receptor (LR)-mediated cell-cell communications for spatially resolved transcriptomic data. Here, we propose SpaCCC to facilitate the inference of spatially resolved cell-cell communications, which relies on our fine-tuned single-cell LLM and functional gene interaction network to embed ligand and receptor genes expressed in interacting individual cells into a unified latent space. The LR pairs with a significant closer distance in latent space are taken to be more likely to interact with each other. After that, the molecular diffusion and permutation test strategies are respectively employed to calculate the communication strength and filter out communications with low specificities. The benchmarked performance of SpaCCC is evaluated on real single-cell spatial transcriptomic datasets with remarkable superiority over other methods. SpaCCC also infers known LR pairs concealed by existing aggregative methods and then identifies communication patterns for specific cell types and their signalling pathways. Furthermore, spaCCC provides various cell-cell communication visualization results at both single-cell and cell type resolution. In summary, spaCCC provides a sophisticated and practical tool allowing researchers to decipher spatially resolved cell-cell communications and related communication patterns and signalling pathways based on spatial transcriptome data.

## 1 Introduction

Multicellular organisms rely on well-organized cell-cell communications (CCCs), which include sender cells, receiver cells, ligand-receptor interactions (LRIs), and the resulting downstream biological responses, to carry out critical biological processes [1]. With prior knowledge of known LRIs and single-cell RNA sequencing (scRNA-seq) data, specific algorithms have been developed to study the cellular communication network by mapping cell-specific LRIs in complex tissues [2, 3]. A common strategy for inferring CCCs involves integrating the abundance of ligands and receptors, taking into account their expression correlation, differential expression, and co-expression (sum, mean, or product) [4–6]. Although scRNA-seq data provide rich details on the genes contributing to CCCs, the loss of spatial information caused by dissociating tissues into single cells limits the usefulness of current algorithms to study CCCs in tissues with spatial structure [7].

Several spatial barcoding and imaging-based methodologies have enabled the characterization of whole or mostly whole gene expression while retaining spatial information in spatially resolved transcriptomics (ST) [8]. This advancement has greatly enhanced different fields of biological and biomedical studies, and has brought the study of cellular communication to a higher level of spatial resolution [9]. Given the fact that intercellular secreted signaling is constrained to space, inferring CCCs based on ST is essential for understanding the cellular tissue functions and disease progression [10].

In recent years, a variety of methods have been proposed to infer CCCs for spatially resolved transcriptomic data. For example, Pham *et al*. [11] presented stLearn, which combines two analysis streams, including ligand-receptor interaction activity and different cell type co-localisation, into one interaction measure to identify hotspots where CCCs are more likely to occur. Dries *et al*. [12] presented Giotto, which utilizes a random permutation strategy of the cell type labels within a defined spatial network to determine the ratio of observed-over-expected frequencies between two cell types. Shao *et al*. [13] presented SpaTalk, which integrated the principles of the ligand-receptor proximity and ligand-receptor-target (LRT) co-expression, and then utilized the knowledge-graph-based method to model and score the LRT signaling network between spatially proximal cells. However, these methods all perform CCC analysis at the level of the cell (spot) type or cluster, discarding single-cell (spot)-level information. Biologically, CCC does not operate at the level of the group; rather, such interactions take place between individual cells. Furthermore, these CCC analyses based on original gene expression profiles are limited by certain technical limitations inherent to scRNA-seq, and thus tend to introduce bias or noise that can affect the accuracy of CCC results [14, 15].

The rapid development of scRNA-seq has generated tens of millions of single-cell data, forming comprehensive data atlases such as the Human Cell Atlas [16]. This allows large language models (LLMs), which have recently achieved unprecedented success in various fields due to their ability to learn from extensive datasets, to be applied to the analysis of single-cell data. Examples of these models include scGPT [17], scBERT [18], Geneformer [19] and so on. In particular, expanding on the self-supervised pre-training in LLMs, scGPT achieves superior performance to multiple downstream tasks such as genetic perturbation prediction, multi-omic integration, cell type annotation and so on. However, to date, it still lacks methods to take advantage of LLMs to infer ligand-receptor (LR)-mediated cell-cell communications for spatially resolved transcriptomic data.

Here, we proposed spaCCC to infer cell-cell communications for spatially resolved transcriptomic data. Specifically, spaCCC first relied on our fine-tuned single-cell LLM and functional gene interaction network to embed ligand and receptor genes expressed in interacting individual cells into a unified latent space. Second, the ligand-receptor pairs with a significant closer distance in latent space were taken to be more likely to interact with each other. Third, molecular diffusion and permutation test strategy were respectively employed to calculate the communication strength and filter out communications with low specificities. In the results, we demonstrated the effectiveness of ligand and receptor embeddings based on the fine-tuned LLM and functional gene interaction network. The benchmarked performance of SpaCCC was evaluated on real single-cell spatial transcriptomic datasets with remarkable superiority over other methods. Furthermore, we also demonstrated that SpaCCC can infer known LR pairs concealed by aggregative methods. Additionally, spaCCC provides the identification of communication patterns for specific cell types and their signalling pathways, and offers a rich suite of visualization options (Circos plot, Heatmap plot, Bubble plot, Network plot, etc.) to present the CCC results with both the single-cell and cell type resolution. In summary, spaCCC provides a sophisticated and practical tool allowing researchers to decipher spatially resolved cell-cell communications and related communication patterns and signaling pathways based on spatial transcriptome data.

## 2 Material and methods

### 2.1 Datasets and data pre-processing

The input of spaCCC is a gene-by-cell count matrix with annotated cell types, a cell position matrix and an histopathological picture. We applied spaCCC to two spatially resolved transcriptomic (ST) dataset on the 10x Visium platform, including human renal cell carcinoma and human breast cancer. The renal cell carcinoma dataset was obtained from the STOmicsDB database (Dataset ID: STDS0000223) [20], which is a comprehensive repository of literature and datasets related to spatial transcriptomics topics. This dataset contains 4510 cells of 5 cell types (**Supplementary Fig. 1.a**). The human breast cancer dataset was obtained from the 10x Genomics website (Visium Demonstration, Human Breast Cancer, Block A Section 1) by the reference [21]. It contains 3798 cells of 9 cell types (**Supplementary Fig. 1.b**). The expression matrix for both dataset were normalized using the LogNormalize method [22]. In addition, the prior knowledge of ligand-receptor interactions in the OmniPath [23] database was adopted. Only ligand-receptor pairs expressed in at least 10% of cells in the respective datasets were considered. On the other hand, the functional gene interaction network was generated by the latest STRING database (version 12.0) [24]. The functionally related signaling pathways for the ligand-receptor interactions were obtained by the systematically curated classification in CellChat [6].

### 2.2 Evaluation metrics

The performance of spaCCC for annotating cell types was evaluated by the standard multi-classification metrics including Accuracy, Precision, Recall, and Macro-F1, which are defined for per cell type *c* as follows:

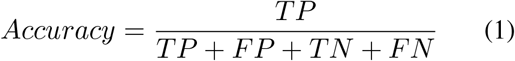

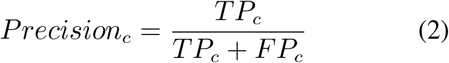

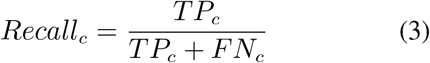

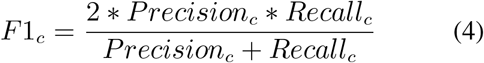

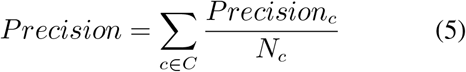

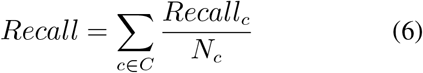

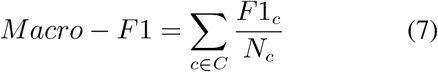

The ligand and receptor embeddings generated by spaCCC was evaluated by the standard binary-classification metrics, additionally including Sensitivity (calculated equal to recall), Specificity, Matthews correlation coefficient (MCC), Receiver Operating Characteristic (ROC) Curve, the areas under the ROC Curve (AUC), Precision Recall (PR) Curve and the areas under the PR Curve (AUPRC), which are defined as follows:

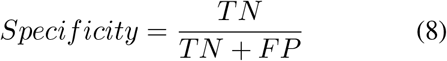

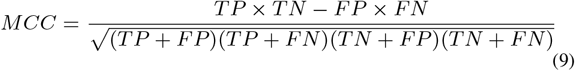

Here, TP, TN, FP, FN respectively means the number of true positives, true negatives, false positives, and false negatives in the prediction results. The ROC and PR curves are commonly used visualization methods in binary classification. They respectively plot the TP rate or Precision against the FP rate or Recall at various discrimination thresholds. The AUC and AUPRC are quantitative evaluation indicators. Their values range between 0 and 1, and the closer the values are to 1, the better is the fit of the model.

On the other hand, the performance for inferring significantly ligand-receptor interactions (LRIs) between spaCCC and different analysis methods was evaluated by the Jaccard index, which is defined by dividing the overlapping LRIs number between two CCC analysis tools by their total LRIs number:

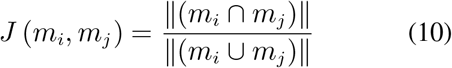

where *m*_*i*_ and *m*_*j*_ respectively denote the inferred LRIs by two different CCC analysis methods, *m*_*i*_ ∩ *m*_*j*_ and *m*_*i*_ ∪ *m*_*j*_ respectively denote the intersection and union of the inferred LRIs, ∥. ∥ denotes the number of the inferred LRIs in the set.

### 2.3 Fine-tuning strategy on pre-trained large language model

By drawing comparisons between language construction and cell biology, we can see that just as texts are comprised of words, cells are defined by genes. As a result, models for single-cell biology are being increasingly developed as foundational frameworks. Among them, scGPT [17], which utilizes a generative pre-trained transformer model that has been trained on a repository of more than 33 million cells, achieves superior performance across diverse downstream applications. Fine-tuning scGPT based on single-cell data, we aim to learn both gene and cell embeddings (similar to word and sentence embeddings in natural language generation) so that genes and cells can be better characterized based on the genes expression in the cells.

Specifically, the gene names and expression values were first preprocessed as input to scGPT. It used gene names as tokens and assigned a unique integer identifier to each gene *g*_*j*_, denoted as 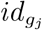. Additionally, it also added special tokens in the token vocabulary, including *< pad >* for padding the input to a fixed length and *< cls >* for aggregating all genes into a cell representation. On this basis, it represented the input gene tokens of each cell *i* using a vector 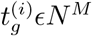:

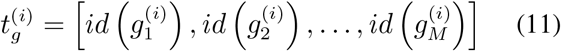

where *N* is the gene numbers and *M* is a pre-defined maximum input length.

Due to variations in sequencing depths and genes that are expressed at low levels, the data scales differ significantly among different sequencing sample batches [25]. Hence, a value binning technique was proposed by scGPT to transform all expression counts into relative values, in order to deal with the scale difference. The raw absolute values were calculated and split into *B* consecutive intervals [*b*_*k*_, *b*_*k*+1_], where *k* ϵ 1, 2, …, *B*. For each non-zero expression count in cell *i*, the binned expression value 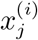 was obtained by:

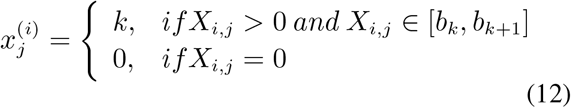

This binning technique ensures that the semantic meaning of 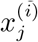 is the same across cells from different sequencing batches. On this basis, the conventional embedding layers *emb*_*g*_ map each gene token to an embedding vector of fixed length *D*. The binned expression values are mapped by the fully connected layers *emb*_*x*_. The final embedding *h*^(*i*)^ for cell *i* is obtained by:

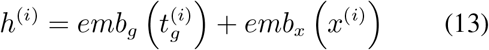

The model was then trained efficiently to capture both gene-gene and gene-cell relation using a generative training strategy with specialized attention masks. To optimize the model for predicting expression values of unknown genes, a gene expression prediction objective was applied:

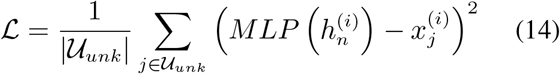

where the multi-layer perceptron is denoted by MLP, the element number of the set is obtained by the operation, the actual gene expression value is denoted by 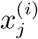, and the set of the output positions for unknown genes is denoted by *U*_*unk*_.

In this work, we used gene expression prediction as fine-tuning objective to obtain the latent embeddings of genes, which contains the interactions between different genes. In particular, a random subset of gene tokens and their corresponding expression values *x*(*i*) were masked for each input cell, and the expression values at the masked positions were accurately predicted by optimizing the pre-trained scGPT. This fine-tuning objective enhanced the model’s ability to encode co-expression patterns among the genes in the dataset, resulting in more effective gene expression analysis. The mean squared error at the masked positions, which are denoted as *M*_*mask*_, was minimized by the objective:

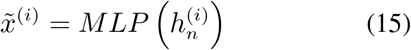

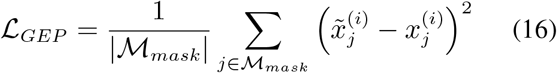

where 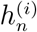 denotes the output of the transformer layer and 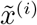 denotes the predicted expression values for cell *i*. Finally, the integrated gene embeddings (Equation 12) were retrieved after model convergence.

### 2.4 Graph embedding strategy on functional gene interaction network

Given the dense and weighted nature of functional gene interaction network, we adopted node2vec+ [26] to extract the latent embeddings of ligands and receptors on the network. This method, an extension of node2vec [27], accommodates edge weights in the calculation of walk biases, resulting in a more accurate representation of ligands and receptors in the low-dimensional space. Specifically, the graph neural networks (GNNs) and graph embedding methods have shown remarkable performance in prediction tasks, particularly in computational biology, such as gene-gene interaction prediction [28], essential protein prediction [29], disease gene prediction [30] and so on. Both GNNs and embedding methods have the same goal of projecting nodes in the graph to a feature space. Among them, the embedding methods are more suitable for our model because GNNs typically require labeled data and initial node features as input. Node2vec is a method for unsupervised node embedding commonly used in different tasks and relies on second-order random walk-based embeddings. However, its effectiveness is diminished when applied to our functional gene interaction due to an inability to recognize out edges. Node2vec+ addresses this limitation by enhancing the second-order random walk approach, specifically tailored for weighted graphs through the consideration of edge weights. Last but not least, it has been demonstrated that node2vec+ outperforms GNNs and node2vec for learning robust node representations in the same STRING networks as ours [26].

### 2.5 Training strategy

In the cell type annotation task, we partitioned 85% of the cell sample in the dataset into the training set and 15% of the cell sample into the test set. We run 50 epochs using the training set to fine-tune the large language model of scGPT on human. The loss function consists of two parts, the mean squared error (mse) loss for mask gene prediction and the cross-entropy loss for cell type annotation. The model with the smallest loss function is used as the final model to predict the test set. On the other hands, in the ligand-receptor interactions prediction task, the gene expression profiles of all cell samples were used to fine-tune the large language model of scGPT on human. The loss function is only the mean squared error (mse) loss for mask gene prediction. The model with the smallest loss function on the epoch (200 in this task) is used to generate the embeddings for ligands and receptors. Both tasks also adopted the Dropout strategy (0.2 in this work), Adam optimizer (initial learning rate of 1e-4 and weight decay of 1e-4) and the StepLR learning rate decay method (gamma of 0.9 and step size of 10).

### 2.6 Determining the statistical significance of ligand-receptor interactions

We considered two genes with a significant closer distance to be more likely interacted. Genes were projected into the hidden space according to the fine-tuning model as well as the interaction network. We first calculated the Euclidean distance *d*_*ij*_ = ∥*E*_*i*_ − *E*_*j*_ ∥ for every pairwise combination of gene *i* and gene *j* in scRNA-seq data, where *E*_*i*_ and *E*_*j*_ means the representations of them in the low dimensional embedding. After that, we implemented a permutation test strategy to identify gene pairs with significant closer distance among all combinations under a null hypothesis, which was constructed by selecting *n* background gene pairs (*n* = 200 in this work) for each gene pair. The average expression levels of the two genes within each background gene pair are similar to those of the corresponding query gene pair. Utilizing the Euclidean distance of these background gene pairs, we derived the null distribution for each gene pair. On this basis, the gene pairs expressed in more than 10% of cells, present in the Ominipath database, and significantly closer than background genes (*p*-value*<*0.05) were determined as the statistical significance of ligand-receptor interactions in this work.

### 2.7 Calculation of the communication strength at single cell resolution

To calculate the communication strength of the ligand-receptor mediated signaling interactions between individual cells, the strategy of HoloNet [21] was adopted in this work. Specifically, the mass action and molecular diffusion in chemistry inspired us to use Gaussian function to model cell-cell communication in the spatial transcriptomic data. On this basis, the communication strength 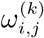 between the ligand *L*_*k*_ expressed on the sender cell *C*_*i*_ and the receptor *R*_*k*_ expressed on the receiver cell *C*_*j*_ can be expressed by the expression level of *L*_*k*_, the expression level of *R*_*k*_ and the spatial distance between *C*_*i*_ and *C*_*j*_:

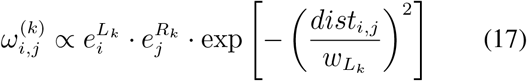

where 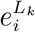 and 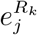 means the expression level of the ligand *L*_*k*_ and receptor *R*_*k*_, and *dist*_*i,j*_ means the Euclidean distance between the sender cell *C*_*i*_ and the receiver cell *C*_*j*_. The ∝ means a proportional correlation and was set to 1 in this work. Furthermore, the diffusion ability of the ligand, denoted as *w*_*L*_*k*, governs the coverage area with a diameter (*d*) of ligands originating from a sender cell (the default setting in Visium datasets establishes *d* as 255*µm*) [31, 32]. It was defined as follows:

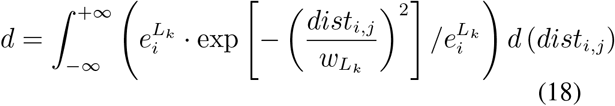

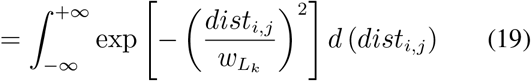

On this basis, 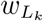 can be denoted as follows:

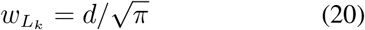

### 2.8 Filtering out ligand-receptor interactions with low specificities

To reserve the ligand-receptor interactions (LRTs) that actively communicating, we used a permutation test strategy developed from the stLearn [11] to calculate the specificity of each ligand-receptor interaction. Specifically, 200 background gene pairs was first randomly selected for each LRT. The two genes within each background gene pair have the most similar average expression levels to the ligand and receptor, respectively. Subsequently, the null distribution for each LRT was generated by calculating the communication strength of each background gene pairs. For a particular LRT, the communication strength of its background gene pairs between cell *i* and *j* is obtained by:

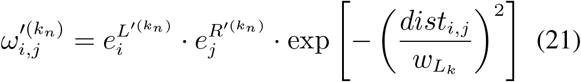

where 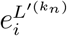 and 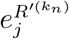 respectively denote the expression level of *k*-th background ligand and receptor in cell *i* and *j*. The communication strength of *n* background gene pairs 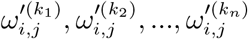 form a null distribution for the particular ligand-receptor pair. Finally, the ligand-receptor interactions whose communication strength is not over 95% of the null distribution will be filtered out.

### 2.9 Ligand-receptor interactions clustering

In this work, the ligand-receptor interactions were clustered by their distribution of spatial communication on the tissue. The ligand-receptor interactions that have similar spatial communication regions were clustered into the same category and vice versa. For two ligand-receptor pairs *u* and *v*, the dissimilarities between them were first calculated:

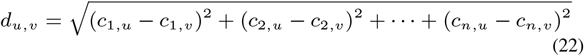

where *c*_*n,u*_ and *c*_*n,v*_ is the eigenvector centrality for cell *n* and ligand-receptor pair *u* and *v*. Subsequently, dissimilarity-based hierarchical clustering was employed to group these interaction pairs. Consequently, we consolidated the communication hotspots within each cluster of ligand-receptor pairs and calculated the cumulative spatial centrality. Furthermore, we employed UMAP dimension reduction to analyze ligand-receptor pairs using the eigenvector centrality matrix and dissimilarity metric.

### 2.10 Calculation of the spatial co-expression of ligand-receptor interactions

In this work, the local bivariate similarity metric was utilized to assess the spatial co-expression between ligands and receptors. This method enabled us to pinpoint the exact location of spatial relationships, and to identify spatial relationships that might occur only in a specific sub-region of our samples. Specifically, the spatially-weighted cosine similarity was utilized in this work, which was natively re-implemented and demonstrated to be the best to identify local ligand-receptor relationships by LIANA+ [33]. For example, the spatially-weighted cosine similarity of spot *i* for a specific ligand-receptor interaction can be defined by performing summation over all spots *n*:

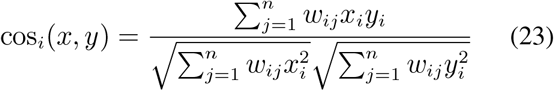

where *x* and *y* denote the horizontal and vertical coordinates of spot *i* on the histological image. *w*_*ij*_ denotes the spatial weights, which are predominantly determined by a set of radial kernels which leverage the inverse Euclidean distance between spots to establish connectivity values ranging from 0 to 1. This approach ensures that spots in close proximity exhibit higher spatial connectivity with each other (assigned a value close to 1), while those considered too distant to interact are assigned a connectivity value of 0.

### 2.11 Visualization of analysis results

Python package scanpy (v1.9.5) [34] was used to generate spatial distributions of cells, genes, spatial co-expressed score and communication hotspots of ligand-receptor interactions on histological images. R package CCPlotR [35] was used to generate the cell-cell comunication results at the level of the cell type. Some other analysis results were visualized by Origin 2023 software.

## 3 Results

### 3.1 Overview of the SpaCCC method

The overview of the workflow for developing SpaCCC is shown in **Fig. 1**. First, the input of spaCCC is common spatial transcriptome data, including single-cell(spot) gene expression profiling, spatial position and cell type information. Second, the statistical significance of ligand-receptor interactions (LRIs) are restricted by fine-tuning on pre-trained large language model (LLM) and graph embedding learning on functional gene interaction network. Finally, the cell-cell communication (CCC) strength at the single-cell and cell-type resolution was calculated (**Fig. 1.a**).

**Fig. 1.**
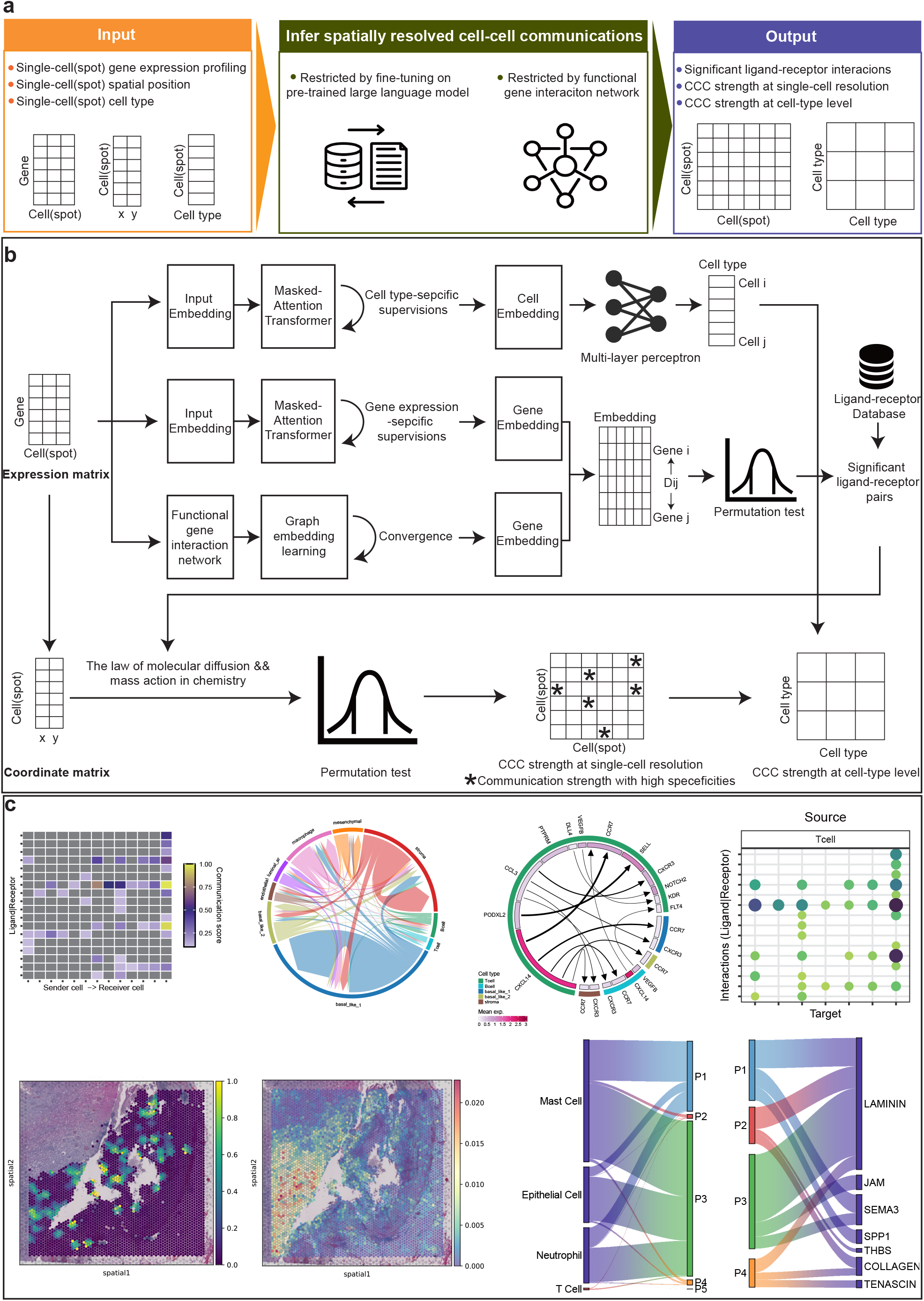
The overview of the workflow for developing spaCCC. **a**. The input, intermediate process for inferring spatially resolved cell-cell communications and output of spaCCC. **b**. The detailed process for determining the statistical significance of ligand-receptor pairs and calculating the cell-cell communication strength at single-cell and cell-type resolution. **c**. Numerous visualization types of spaCCC to decipher the spatially resolved cell-cell communications **(see section 3.1 for details)**.

Specifically, the single-cell(spot) gene expression profiling was first preprocessed as input embedding for fine-turning the scGPT, which is the single-cell pre-trained large language model based on the masked-attention transformer **(section 2.3)**. Based on the cell type-specific and gene expression-specific supervisions, the cells and genes were respectively represented as embedding vectors. The cell embeddings were further used to train and predict the cell types by multi-layer perceptron. On the other hand, the genes in the expression profiling were constructed as a functional gene interaction network. The second type of gene embeddings were obtained by using graph embedding learning method Node2vec+ **(section 2.4)**. Based on the two types of gene embeddings, we calculated the Euclidean distance under the latent hidden space for each gene pairs, followed by determining the gene pairs that were significant closer using the permutation test. The gene pairs that significantly closer in the latent hidden space and present in the priori ligand-receptor database OminiPath were identified as the statistical significance of ligand-receptor interactions **(section 2.6)**. After that, the CCC strength at single cell resolution was calculated using the law of molecular diffusion and mass action in chemistry based on the single-cell(spot) gene expression profiling and significant ligand-receptor pairs **(section 2.7)**. To reserve the ligand-receptor interactions that actively communicating, spaCCC filtered out the interactions with low specificities **(section 2.8)**. Finally, spaCCC also provides the CCC results at the cell-typen level by summing the results of single-cell resolution to meet the needs of some researchers (**Fig. 1.b**).

SpaCCC also includes numerous visualization types to decipher the spatially resolved cell-cell communications, such as the heatmap of the CCC strength at single-cell resolution, where each row represents a LRI pair and the elements in each column represent the CCC strength from a sender cell to a receiver cell. The chord plot of CCC strength between each pair of cell types, where different colors represent different cell types, the edge thickness is proportional to the communication strength, and edge direction goes from the sender cell type to the receiver cell type. The circos plot of communication strength of LRI pairs between different cell types, where the colors of the outer ring represent different cell types, the color shade of the inner ring represents the mean expression of the genes in the corresponding cell types, and the thickness of the arrow is proportional to the communication strength, with the direction of the arrow goes from the ligand to the receptor. The dot plot of communication strength of LRI pairs from a sender cell type to a recevier cell type. Furthermore, the umap plot of the spatially co-expressed score of LRI pairs, the umap plot of the communication pattern of LRI pairs, and the sankey plot of the communication pattern and signaling pathway of LRI pairs were also included in spaCCC (**section 2.11, Fig. 1.c**).

### 3.2 Performance evalution of fine-tuning strategy on pre-trained large language model

The fine-tuning strategy on pre-trained large language model by spaCCC is the foundation for subsequent analyses. To evaluate its performance, two spatially resolved transcriptomic (ST) dataset, including human renal cell carcinoma and human breast cancer (**Supplementary Fig. 1**), and two different tasks, including cell type annotation and ligand-receptor interactions (LRIs) prediction, were selected. First, we evaluated the performance of the fine-tuned model on cell type annotation for our two dataset. The dataset was evenly divided into five parts: four parts were used as training data for fine-tuning and one part as test data for evalution. We visualized the spatial distribution of real and predicted cell types for the two test data (**Fig. 2.a and c**), which was highly consistent in that region. We used heatmap to further visualize the confusion matrix between real and predicted cell types,with darker colors indicating higher concordance and lighter colors indicating lower concordance (**Fig. 2.b and d**). Dark colors on the diagonal indicated that the predicted cell types were frequently correct, only except for rare cell types with extremely low cell numbers in the reference partition. Furthermore, the annotation quality was also evaluated quantitatively by accuracy, precision, recall, and macro F1-score metrics (**Supplementary Table 1**). The annotation accuracy of both dataset was greater than 80%. For renal cell carcinoma with fewer cell types, the accuracy even reached 96.2%. These results suggested that the fine-tuning strategy on pre-trained large language model was able to effectively learn cell representations composed of latent embeddings of genes.

**Fig. 2.**
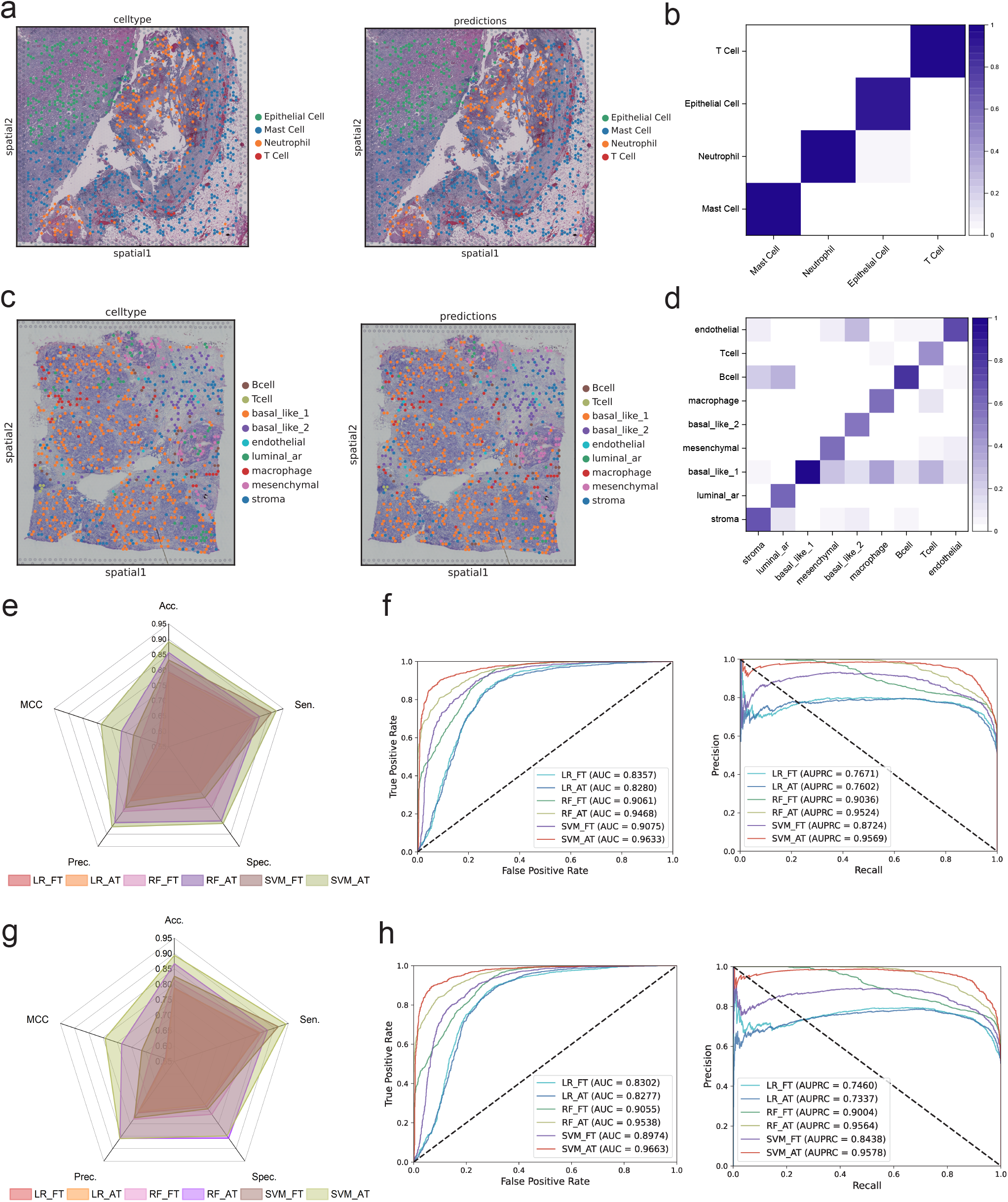
Performance evalution of fine-tuning strategy on pre-trained large language model using two spatially resolved transcriptomic datasets and two different tasks. **a**. The spatial distribution of real and predicted cell types for renal cell carcinoma dataset. **b**. The confusion matrix between real and predicted cell types for renal cell carcinoma dataset. **c**. The spatial distribution of real and predicted cell types for human breast cancer dataset. **d**. The confusion matrix between real and predicted cell types for human breast cancer dataset. **e**. The radar chart of five common metrics (Accuracy: Acc., Precision: Prec., Sensitivity: Sen., Specificity: Spec., Matthews Correlation Coefficient: MCC) under 5-fold cross-validation in the ligand-receptor interaction prediction task for renal cell carcinoma dataset. **f**. The receiver operating characteristic (ROC) curves, precision-recall (PR) curves, the area under the ROC curve (AUC) values, and the area under the PR curve (AUPRC) values under 5-fold cross-validation in the ligand-receptor interaction prediction task for renal cell carcinoma dataset. **g**. The radar chart for human breast cancer dataset. **h**. The ROC curves, PR curves, the AUC, and the AUPRC values for human breast cancer dataset.

To more accurately evaluate the performance of LRIs prediction task for demonstrating the validity of ligand and receptor gene embedding, which was the foundation of our downstream CCC inference, a 5-fold cross-validation method was utilized. Specifically, we used the LRI paris inferred from each dataset as the positive sample and an equal number of randomly generated LRI pairs (positive sample excluded) as the negative sample. The training and test data were divided in the same way as the cell type annotation task. The above measurement was repeated 5 times and the average value was taken as the final result. We simultaneously evaluated both types of ligand and receptor gene embedding, which respectively obtained from functional gene interaction network (AT) and the fine-tuned large language model (FT). Different classification models (Logistic Regression: LR, Random Forest: RF, Support Vector Machine: SVM) and five common metrics (Accuracy: Acc., Precision: Prec., Sensitivity: Sen., Specificity: Spec., Matthews Correlation Coefficient: MCC) were utilized for this evaluation. We visualized the above metrics by using radar charts (**Fig. 2.e and g**), and the detailed results were also presented in **Supplementary Table 2 and 3**. Furthermore, we plotted the receiver operating characteristic (ROC) and precision-recall (PR) curves, and calculated the area under the ROC curve (AUC) and the area under the PR curve (AUPRC) values (**Fig. 2.f and h**). For the renal cell carcinoma dataset, the LRI prediction with the functional gene interaction network-based embeddings of ligands and receptors achieved the highest average AUC and AUPRC value of 0.9633 and 0.9569 under 5-fold cross-validation. On the other hand, the fine-tuned large language model-based embedding achieved the highest average AUC and AUPRC value of 0.9075 and 0.9036, respectively. For the human breast cancer dataset, four values above was 0.9663, 0.9578, 0.9075 and 0.9036, respectively. These excellent results suggested that the both types of embeddings extracted by spaCCC were able to effectively learn representations of ligands and receptors, which made the subsequent analyses accurate and reliable.

### 3.3 Comparison of spaCCC with other existing methods

The first part of analyzing cell-cell communications is to infer significantly ligand-receptor (LR) pairs from transcriptomics. We compared the performance of spaCCC for inferring LR pairs with 7 common existing methods (SingleCellSignalR [5], NATMI [36], CellPhoneDB [4], Connectome [37], CellChat [6], along with logFC [33] and a geometric mean [33]) on the renal cell carcinoma and human breast cancer dataset. In the experiment, all methods relied on the same LR resources in the OmniPath [23] database and considered only the ligands and receptors expressed in more than 10% cells, refer to CellPhoneDB method. The following results parameters were used to select the top 1000 LR pairs from each method for comparision: LRscore (SingleCellSignalR), spec weight (NATMI), cellphone pvals (CellPhoneDB), scaled weight (Connectome), cellchat pvals (CellChat), lr logfc (logFC), gmean pvals (geometric mean) and spaccc pvals (spaCCC).

First, we looked at the overlap between the 1000 highest ranked LR interactions inferred by each method. The Jaccard index, defined as the ratio of the intersection to the union of sets, was utilized to estimate the relative agreement between these methods. **Fig. 3.a and c** respectively showed the comparison results of renal cell carcinoma and human breast cancer dataset. The heatmaps indicated pairwise similarity according to the Jaccard index (darker being stronger agreement). The histograms indicated the sum of the Jaccard index for each method versus the other methods. By considering both datasets together, we found that the overlap of the top inferred interactions was consistently low when using any different methods. One possible reason for the discrepancy is that the existing methods are based on the same original gene expression but different screening strategies. spaCCC has the third best-Jaccard index, behind the logFC and NATMI methods. However, by further observing the heatmaps, the Jaccard index of spaCCC had a more evenly distribution on both datasets. In contrast, all other methods, including logFC and NATMI, will more or less have extremely low overlap with one or more methods. The experimental results demostrated that spaCCC was able to obtain more stable and robust LR inference results than other existing methods by utilizing their latent embeddings.

**Fig. 3.**
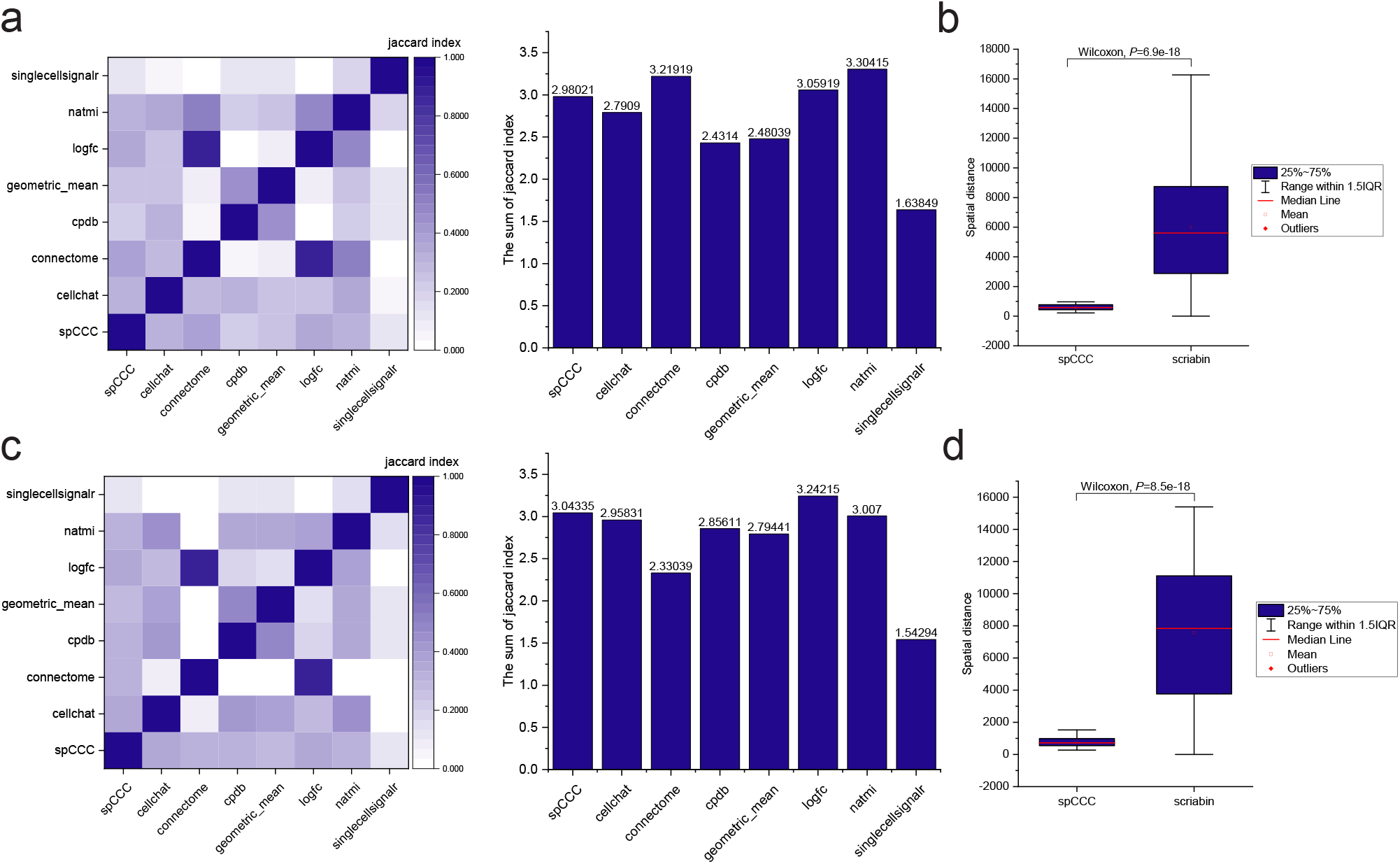
Comparison of spaCCC with other existing methods. **a** The heatmap indicated pairwise similarity between the 1000 highest ranked LR interactions inferred by each method on renal cell carcinoma dataset (darker being stronger similarity). The Jaccard index was defined as the ratio of the intersection to the union of sets. The histogram indicated the sum of the Jaccard index for each method versus the other methods. **b** The Box plot indicated the comparison results of spaCCC and Scriabin for the spatial distance distribution of the top 1000 cell-cell pairs with the highest communication strength on renal cell carcinoma dataset. Shown is an exact twosided *P* value from the Wilcoxon rank-sum test. **c** The heatmap indicated pairwise similarity between the 1000 highest ranked LR interactions inferred for every method on human breast cancer dataset (darker being stronger similarity). The Jaccard index was defined as the ratio of the intersection to the union of sets. The histograms indicated the sum of the Jaccard index for each method versus the other methods. **d** The Box plot indicated the comparison results of spaCCC and Scriabin for the spatial distance distribution of the top 1000 cell-cell pairs with the highest communication strength on human breast cancer dataset. Shown is an exact twosided *P* value from the Wilcoxon rank-sum test.

Furthermore, we also looked at the spatial distance distribution of the top 1000 cell-cell pairs with the highest communication strength. The distribution of spatial distance could support the rationality of cell-cell communication analysis results from the side. A biologically relevant hypothesis was that the strength of cell-cell communications becomes weaker with increasing spatial distance. As all the above methods performed differential CCC analyses by aggregating data at the level of cell type or cluster. We further performed comparisons using Scriabin with its default parameters [38], a flexible and scalable framework for comparative analysis of cell-cell communications at single-cell resolution. Box plots shown the comparison results of spaCCC and Scriabin. An exact twosided *P* values were calculated using the Wilcoxon rank-sum test. **Fig. 3.b and d** shown that in the results of spaCCC at single-cell resolution, the spatial distance of cell-cell pairs with greater communication strength is significantly closer than Scriabin on both datasets, which demostrated that the inference results of spaCCC were more consistent with actual biological phenomena.

### 3.4 spaCCC infers known LR pairs concealed by aggregative methods and then filters non-specific CCCs

Exsiting methods for inferring significantly LR pairs from transcriptomics generally function by aggregating ligand and receptor expression values for groups of cells to further infer which groups of cells are likely to interact with one another [39], such as CellPhoneDB [4], Connectome [37], CellChat [6], et al. These aggregative methods could introduce a bias in the inference results, favoring ligands that are specifically high expressed in the sender cell type and receptors that are specifically high expressed in the receiver cell type. As a result, LR pairs with low expression levels may be concealed. To demonstrate this, we selected the most widely employed CellPhoneDB method to infer LR pairs from our renal cell carcinoma dataset. Taking neutrophils as sender cells and the other three cell types as receiver cells, we performed a statistical analysis on the average expression of the LR pairs inferred by CellPhoneDB in each cell type. From **Fig. 4.a**, we can observe that the ligands and receptors, as inferred by CellPhoneDB, exhibit relatively high expression levels in the respective sender and receiver cells. Furthermore, we also analyzed the inference results of the recently published Connectome method from our human breast cancer dataset. Taking Tcell as sender cells, and macrophage and Bcell as receiver cells, we selected the top 10 LR pairs based on its scaled weight parameter for observation. From **Supplementary Fig. 2.a and b**, the ligands and receptors, as inferred by Connectome, also exhibit relatively high expression levels in the respective sender and receiver cells. These experimental results highlight the limitations of the current aggregative methods, which tends to prioritize highly expressed LR pairs in the respective sender and receiver cells, may overlooking the potential communications from lowly expressed ones.

**Fig. 4.**
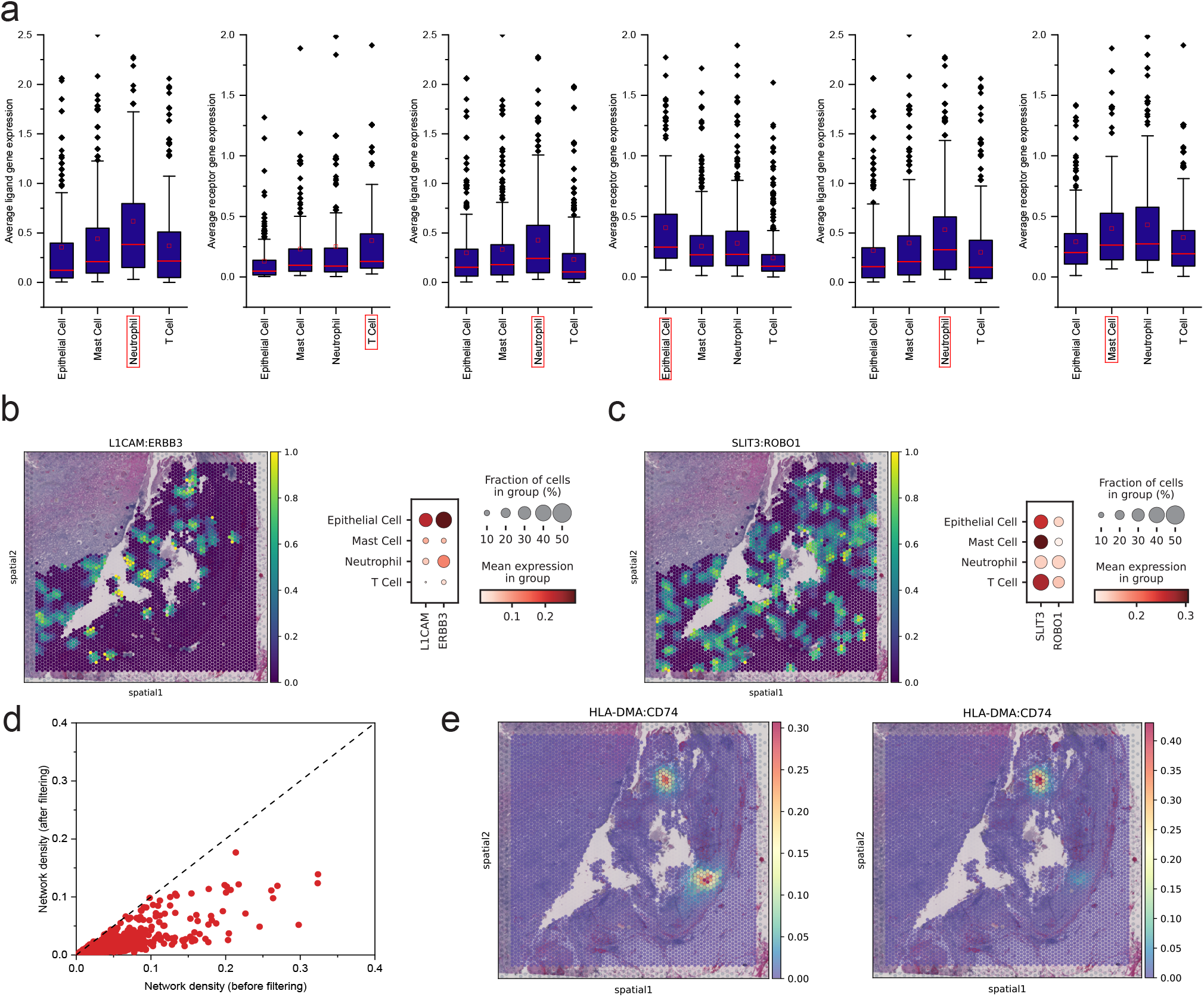
Based on the renal cell carcinoma dataset, spaCCC infers known LR pairs concealed by aggregative methods and then filters non-specific cell-cell interactions. **a**. A statistical analysis on the average expression of the LR pairs inferred by CellPhoneDB in each cell type (Sender cell: neutrophils; Receiver cell: TCell, Epithelial Cell and Mast Cell). **b**. The umap plot of the spatially co-expressed score of L1CAM:ERBB3 pairs in neutrophil and mast cells calculated by using LIANA+. The dot plot of ligand (L1CAM) and receptor (ERBB3) expressed in neutrophils and mast cells. **c**. The umap plot of the spatially co-expressed score of SLIT3:ROBO1 pairs in neutrophil and mast cells calculated by using LIANA+. The dot plot of ligand (SLIT3) and receptor (ROBO1) expressed in neutrophils and mast cells. **d**. The changes of network density before and after filtering non-specific cell-cell interactions by spaCCC for each LR pair. **e**. The umap plot of the communication hotspots of HLA-DMA:CD74 pair before and after filtering non-specific cell-cell interactions by spaCCC.

Correspondingly, spaCCC does not have this limitations of aggregative methods. Instead of inferring directly from the original gene expression, spaCCC utilizes the latent embeddings from two views that allows it not to conceale the low expression LR pairs. For example, two ligand:receptor pairs, L1CAM:ERBB3 and SLIT3:ROBO1, were inferred by spaCCC between neutrophil and mast cell, while CellPhoneDB concealed them. From the dotplot in **Fig. 4.b and c**, we can observe that these two LR pairs were not significantly highly expressed in neutrophil and mast cell. On this basis, we calculated the spatially co-expressed score of them in neutrophil and mast cells by using LIANA+ [33]. The spatial weights in LIANA+ are by default defined as a family of radial kernels that use the inverse Euclidean distance between spots to bind the weights between 0 and 1, with spots that are closest having the highest spatial connectivity to one another, while those that are thought to be too far to be in contact are assigned 0. As shown in the umap plot of **Fig. 4.b and c**, two LR pairs were highly co-expressed at multiple locations in the space occupied by neutrophil and mast cells. This effectively means that these ligands and receptors are most likely communicating at these locations. Similarly, we also calculated the spatially co-expressed score of four LR pairs in our human breast cancer dataset (**Supplementary Fig. 2.c and d**), which were inferred by spacCC but concealed by Connectome between Tcell and macrophage, as well as Tcell and Bcell (**Supplementary Fig. 3.a and b**). These experimental results highlight that the strategy of spaCCC by using latent embeddings could infer known LR pairs concealed by current aggregative methods.

Based on the inferred LR results, spaCCC provided the cell-cell interaction results for each LR pair to analyze communication for each cell(spot)-cell(spot) pair in the dataset. However, non-specific widely expressed cell-cell interactions with each LR pair will affect the results of downstream intercellar communication analysis. To determine which cell pairs were actively communicating, spaCCC filtered the non-specific cell-cell interactions **see section 2.8 for details. Fig. 4.d** displays the results before and after filtering non-specific cell-cell interactions for each LR pair in the renal cell carcinoma dataset. Each red point in the figure represents the cell-cell interaction network density for each LR pair, which are distributed under the diagonal. It suggests that spaCCC removes non-specific cell-cell interactions thus reducing the network density. In addition, we visualized the communication hotspots of one of the LR pairs (HLA-DMA:CD74) using the HoloNet [21] method (**Fig. 4.e**). After filtering non-specific cell-cell interactions by spaCCC, the hotspot region of LR communications is more prominent. Similarly, we demonstrated this again on the human breast cancer dataset (**Supplementary Fig. 2.e and f**).

### 3.5 spaCCC identifies communication patterns and corresponding signaling pathways

To further interprete the complex intercellular communication networks, an important question is how multiple cell groups and signaling pathways coordinate to function. We showcase the functionality of spaCCC to identify communication patterns by applying it to both renal cell carcinoma and human breast cancer dataset. **Fig. 5** presents the analysis results of the renal cell carcinoma dataset. **Fig. 5.a** shows the pattern clustering results for ligand-receptor (LR) pairs. Most of the LR pairs are classified as Pattern3 and Pattern1, followed by Pattern4, Pattern2 and Pattern5. In this section, spaCCC clustered the inferred LR pairs into 5 patterns (user-definable hyperparameter) using the hierarchical clustering algorithm relied on the eigenvector centrality vectors of the intercellular communication network corresponding to each LR pair.

**Fig. 5.**
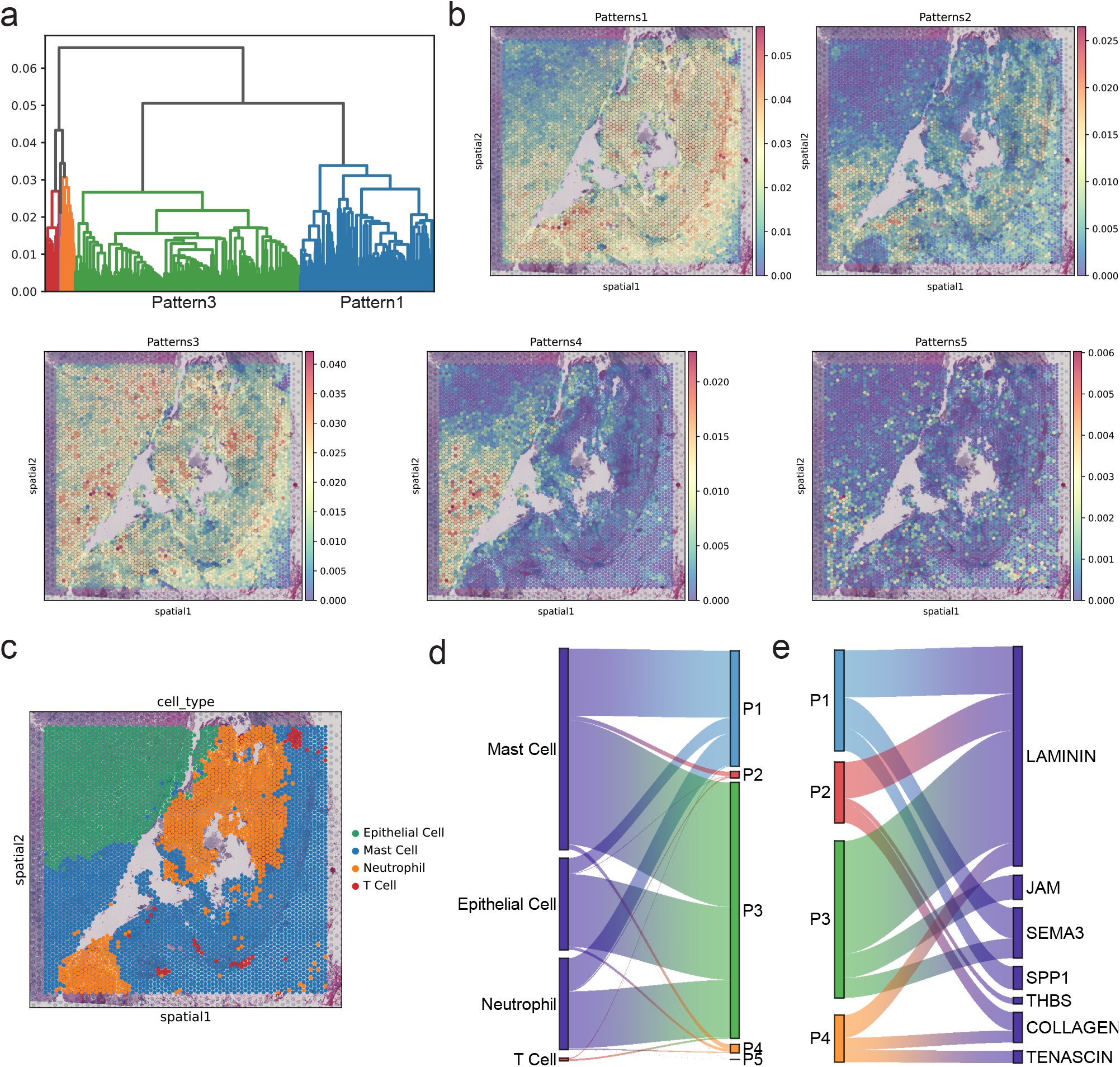
spaCCC employs a pattern recognition method based on the eigenvector centrality to identify communication patterns, as well as the signaling pathways in different cell types of renal cell carcinoma dataset. **a**. Dendrogram for hierarchically clustering all LR pairs. LR pairs were clustered into 5 patterns (user-definable hyperparameter) using a hierarchical clustering algorithm based on the eigenvector centrality vectors of the intercellular communication network corresponding to each LR pair. **b**. The umap plot of the general spatial communication hotspot for each pattern by combining the spatial communication hotspots for all LR pairs within the pattern. **c**. The umap plot of the spatial distribution for each cell type in the dataset. **d**. The sankey plot of the communication strength of cell types in different communication patterns, which are proportional to edge width. **e**. The sankey plot of the proportion of different signaling pathways in each pattern, which are proportional to edge width.

**Fig. 5.b** visualizes the general spatial communication hotspot for each pattern by combining the spatial communication hotspots for all LR pairs within the pattern. **Fig. 5.c** shows the spatial distribution for each cell type in the dataset. Furthermore, spaCCC calculated the communication strength of multiple cell types in each pattern by summing the communication strength at single-cell resolution and classified each LR pair into one of the 229 functionally related signaling pathways based on the CellChat database [6]. By counting the number of signaling pathways in each pattern, spaCCC interpreted the major signaling pathways belonging to them. The sankey plots of **Fig. 5.d and e** respectively show the communication strength of cell types in different communication patterns and the proportions of different signaling pathways in these patterns, which are proportional to edge width.

These visualization types help researchers reveal the major communication patterns between different cell types, the spatial locations where the communication patterns are located and further the major signaling pathways for these patterns. For example, the communication signals between neutrophil and mast cells located on the right side of the spatial distribution are dominated by Pattern1 and Pattern3, which includes multiple pathways, including but not limited to LAMININ and SEMA3. The mast cells located in the lower left corner has a distinct internal communication within them (Pattern4), which also includes multiple pathways, including LAMININ, COLLAGEN and TENASCIN. Furthermore, **Supplementary Fig. 4** presents the analysis results of the human breast cancer dataset. Collectively, spaCCC can identify key features of intercellular communications and their corresponding signaling pathways within a given spatially resolved transcriptomic dataset.

### 3.6 spaCCC provides CCC results at the level of the cell type based on single-cell-level information

To infer which types of cells are likely to interact with one another, existing methods usually work by aggregating ligand and receptor expression values for cell types. But CCC does not operate at the group level biologically; instead, such interactions occur between individual cells. Considering the needs of some researchers, spaCCC also provides CCC results at the level of the cell type by summarizing cellular communication results at single-cell resolution. The communication strength between a pair of cell types is the sum of the communication strengths generated through all LR pairs between individual cells of the belonging cell types. spaCCC also provides a variety of visualization options to further interprete the complex intercellular communication networks. Taking the human breast cancer dataset as an example, **Fig. 6** shows the visualization results. Specifically, the chord plot of **Fig. 6.a** shows the CCC strength between each pair of cell types, where different colors represent different cell types, the edge thickness is proportional to the communication strength, and edge direction goes from the sender cell type to the receiver cell type. The network plot of **Fig. 6.b** shows communication strength between a certain cell type (Tcell as an example) as sender cell and other cell types as receiver cells. The edge thickness is also proportional to the communication strength, and edge direction goes from the sender cell type to the receiver cell type. The heatmap of **Fig. 6.c** shows the CCC strength at single-cell resolution, where each row represents a LRI pair and the elements in each column represent the CCC strength from a sender cell to a receiver cell. The dot plot of **Fig. 6.d** shows communication strength of some ligand-receptor pairs between a certain cell type (Tcell as an example) as sender cell and other cell types as receiver cells. The circos plot of **Fig. 6.e** further shows the communication strength of LRI pairs between different cell types, where the colors of the outer ring represent different cell types, the color shade of the inner ring represents the mean expression of the genes in the corresponding cell types, and the thickness of the arrow is proportional to the communication strength, with the direction of the arrow goes from the ligand to the receptor. The network plot of **Fig. 6.f** and the sigmoid plot of **Fig. 6.g** also shows the ligand-receptor interactions between different cell types. The arrow plot of **Fig. 6.h** shows the ligand-receptor interactions between a certain sender cell type (Tcell as an example) and recevier cell type (basal like 1 as an example). The edge width is proportional to the communication strength. The color shade is proportional to mean gene expression in the cell type. For convenience, only a portion of the ligand-receptor pairs are shown in the above visualization results. The user can specify the number of LR pairs and cell types to be displayed.

**Fig. 6.**
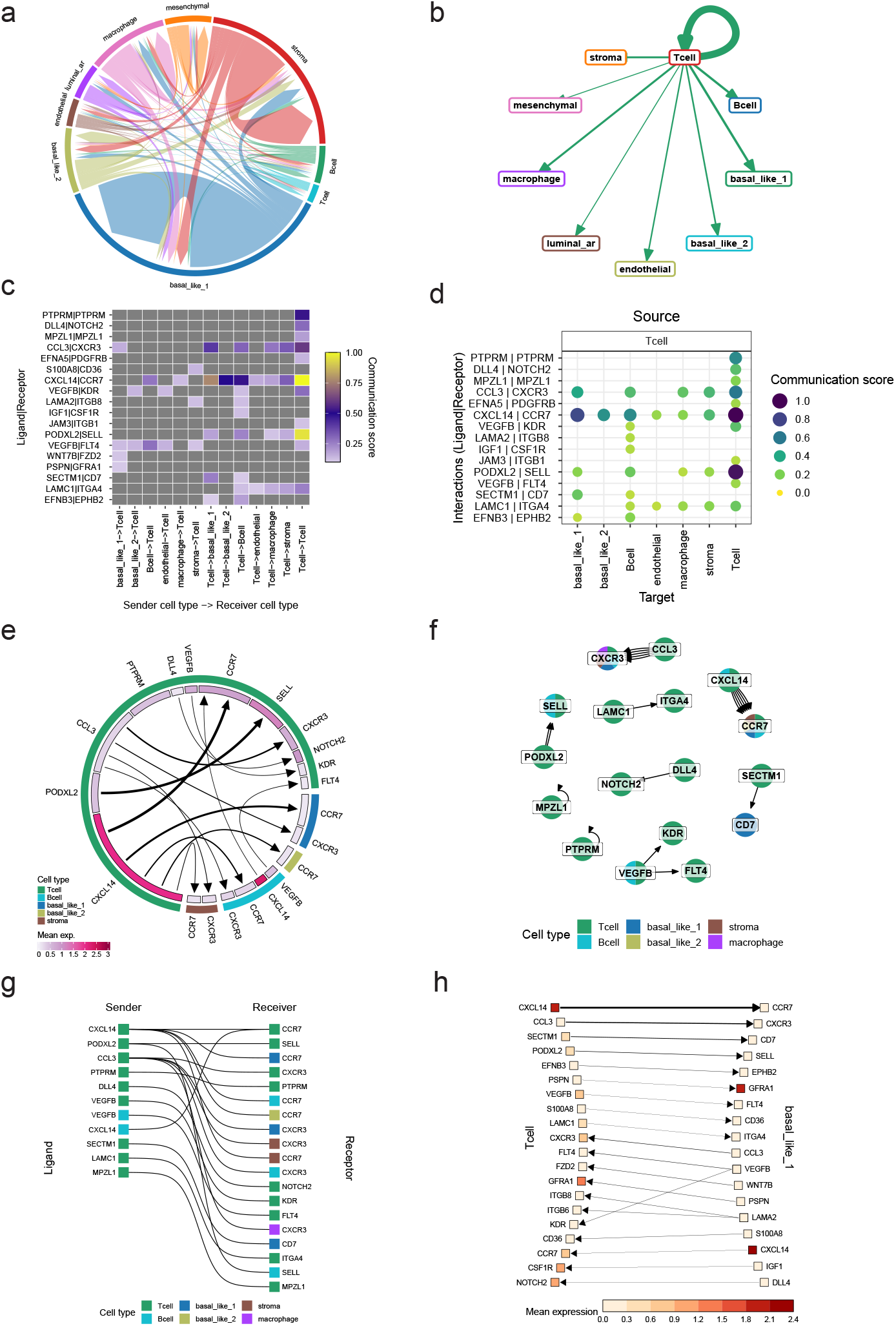
spaCCC provides CCC results at the level of the cell type based on single-cell-level information. **a**. The chord plot shows the CCC strength between each pair of cell types, where different colors represent different cell types, the edge thickness is proportional to the communication strength, and edge direction goes from the sender cell type to the receiver cell type. **b**. The network plot shows communication strength between a certain cell type (Tcell as an example) as sender cell and other cell types as receiver cells. The edge thickness is also proportional to the communication strength, and edge direction goes from the sender cell type to the receiver cell type. **c**. The heatmap shows the CCC strength at single-cell resolution, where each row represents a LRI pair and the elements in each column represent the CCC strength from a sender cell to a receiver cell. For convenience, only a portion of the ligand-receptor pairs are shown. **d**. The dot plot shows communication strength of some ligand-receptor pairs between a certain cell type (Tcell as an example) as sender cell and other cell types as receiver cells. For convenience, only a portion of the ligand-receptor pairs are shown. **e**. The circos plot shows the communication strength of LRI pairs between different cell types, where the colors of the outer ring represent different cell types, the color shade of the inner ring represents the mean expression of the genes in the corresponding cell types, and the thickness of the arrow is proportional to the communication strength, with the direction of the arrow goes from the ligand to the receptor. Mean exp.: the mean expression of the ligand and receptor genes in each cell type. **f**. The network plot of ligand-receptor pairs between different cell types. **g**. The sigmoid plot shows the connection between the ligands in the sender cells and the receptors in the receiver cells. For convenience, only a portion of the ligand-receptor pairs are shown. **h**. The arrow plot shows the ligand-receptor interactions between a certain sender cell type (Tcell as an example) and recevier cell type (basal like 1 as an example). The edge width is proportional to the communication strength. The color shade is proportional to mean gene expression in the cell type. For convenience, only a portion of the ligand-receptor pairs are shown in the above visualization results. The user can specify the number of LR pairs and cell types to be displayed.

## 4 Conclusion

Multicellular organisms rely on cell-cell communications (CCCs) to coordinate the behaviour of individual cells, which enables their differentiation and hierarchical organization. Caused by dissociating tissues into single cells, the single-cell RNA sequencing (scRNA-seq) data lacked spatial information, thus limiting the usefulness of current algorithms to study CCCs in tissues with spatial structure. Given the fact that intercellular secreted signaling is constrained to space, inferring CCCs based on spatially resolved transcriptomic (ST) data is essential for understanding the cellular tissue functions and disease progression. However, the exsiting methods for inferring CCCs based on the ST data either perform at the level of the cell (spot) type or cluster, discarding single-cell(spot)-level information, or perform based on original gene expression, introducing bias that affect the accuracy of the analysis results. Here, we proposed spaCCC to infer cell-cell communications for spatially resolved transcriptomic data, which relies on fine-tuned single-cell large language model and functional gene interaction networks to embed ligand and receptor genes expressed in interacting individual cells into a unified latent space. Based on two real ST datasets, we demostrated that the both types of embeddings extracted by spaCCC were able to effectively learn representations of ligands and receptors, which made the subsequent analyses accurate and reliable. We also demonstrated that the inference results of spaCCC were more stable, robust, and consistent with actual biological phenomena by comparing with existing methods. Moreover, spaCCC could infer konwn ligand-receptor pairs concealed by existing aggregative methods and filter non-specific CCCs. To further interprete the complex intercellular communication networks, spaCCC also provided the identification of communication patterns and their corresponding signaling pathways. Considering the needs of some researchers, spaCCC also provides CCC visualization results at the level of the cell type by summarizing cellular communication results at single-cell resolution. In summary, spaCCC provides a sophisticated and practical tool allowing researchers to decipher spatially resolved cell-cell communications and related communication patterns and signaling pathways based on spatial transcriptome data.

## Acknowledgment

This work was supported by NSFC-FDCT Grants 62361166662; National Key R&D Program of China 2023YFC3503400, 2022YFC3400400; Key R&D Program of Hunan Province 2023GK2004, 2023SK2059, 2023SK2060; Top 10 Technical Key Project in Hunan Province 2023GK1010; Key Technologies R&D Program of Guangdong Province (2023B1111030004 to FFH). The Funds of State Key Laboratory of Chemo/Biosensing and Chemometrics, the National Supercomputing Center in Changsha (http://nscc.hnu.edu.cn/), and Peng Cheng Lab.

